# *De Novo* Phosphoinositide Synthesis in Zebrafish Is Required for Triad Formation but Not Essential for Myogenesis

**DOI:** 10.1101/2020.03.24.005306

**Authors:** Lindsay Smith, Lacramioara Fabian, Almundher Al-Maawali, Ramil R. Noche, James J. Dowling

## Abstract

Phosphoinositides (PIPs) and their regulatory enzymes are key players in many cellular processes and are required for aspects of vertebrate development. Dysregulated PIP metabolism has been implicated in several human diseases, including a subset of skeletal myopathies that feature structural defects in the triad. The role of PIPs in skeletal muscle formation, and particularly triad biogenesis, has yet to be determined. CDP-diacylglycerol-inositol 3-phosphatidyltransferase (CDIPT) catalyzes the formation of phosphatidylinositol, which is the base of all PIP species. Loss of CDIPT should, in theory, result in the failure to produce PIPs, and thus provide a strategy for establishing the requirement for PIPs during embryogenesis. In this study, we generated *cdipt* mutant zebrafish and determined the impact on skeletal myogenesis. Analysis of *cdipt* mutant muscle revealed no apparent global effect on early muscle development. However, small but significant defects were observed in triad size, with T-tubule area, inter terminal cisternae distance and gap width being smaller in *cdipt* mutants. This was associated with a decrease in motor performance. Overall, these data suggest that myogenesis in zebrafish does not require *de novo* PIP synthesis but does implicate a role for CDIPT in triad formation.

## Introduction

The primary function of skeletal muscle is to produce the force that initiates and controls movement. Muscle has a number of unique substructures that are dedicated to force production, including the sarcomere, the neuromuscular junction (NMJ) and the triad (Dowling et al., 2014). As our understanding of the molecular basis of human muscle diseases grows, it is becoming more apparent that many myopathies involve alterations to at least one of these structures (Dowling et al., 2014; Gonorazky et al., 2018; Nance et al., 2012). Of increasing significance are the abnormalities in the structure and function of the triad, which represents the apposition of the T-tubules and the terminal cisternae of the sarcoplasmic reticulum (SR). The key role of the triad is to mediate excitation-contraction coupling (EC coupling), the process by which skeletal muscle translates neuronal signals into muscle contraction (Al-Qusairi and Laporte, 2011; Jungbluth et al., 2018).

Although triad malformations are considered the major cause of muscle weakness in many myopathies (Dowling et al., 2014), the factors that govern the development and maintenance of the triad remain unclear. Recent data has suggested that phosphoinositides may play an important role in triad formation and/or maintenance. Phosphoinositides (PIPs) are a family of membrane phospholipids involved in many essential cell functions, including cellular signaling, endocytosis, and autophagy, and are present in almost all cell types across eukaryotic species (De Camilli et al., 1996) (Balla, 2013). Formation and turnover of the various PIP species are catalyzed by evolutionarily conserved families of kinases and phosphatases (De Matteis and Godi, 2004; Viaud et al., 2016). Dysregulation of PIPs and their metabolic enzymes have been implicated in a number of human diseases, such as congenital myopathies, Charcot-Marie-Tooth Disease (CMT), Alzheimer’s disease, and some forms of cancer (Bunney and Katan, 2010; Lo Vasco, 2018; Nicholson et al., 2011; Volpatti et al., 2019). The consideration of a potential role for PIPs in muscle development comes from two areas of study. One is the work surrounding BIN1, a BAR domain-containing protein that is known to recognize and induce membrane curvature (Peter et al., 2004); (Frost et al., 2009). BIN1 has a PIP-binding domain that interacts with PIP2 (one of the seven PIP sub-species), and this interaction plays a critical role in the formation of T-tubules. Recessive mutations in *BIN1* result in centronuclear myopathy, a severe congenital muscle disease featuring abnormal muscle structure including disturbance of the T-tubule and the triad as a whole. The second line of evidence comes from another form of centronuclear myopathy called X-linked myotubular myopathy or XLMTM (Dowling et al., 2009) (Amoasii et al., 2012). XLMTM is caused by mutations in the PIP phosphatase myotubularin. Mutation in myotubularin causes accumulation of PI3P and leads to abnormalities in the appearance and number of the triad.

In this study, we investigated the role of PIPs in skeletal muscle triad development using the zebrafish model system. Zebrafish is an elegant model for studying skeletal muscle development (Gibbs et al., 2013). Skeletal muscle develops rapidly in zebrafish, muscle fibers are already developing by 24 hours post-fertilization (hpf), with elongated fibers visible by 2 days post-fertilization (dpf). Skeletal muscle is highly prominent in embryos and larvae, and the transparency of developing fish allows muscle fibers to be easily observed. Additionally, zebrafish muscle shares many structural and histological features with mammalian muscle.

To determine the overall requirement for PIPs in muscle development we used the CRISPR/Cas9 technology to generate a *cdipt* zebrafish mutant. CDIPT, also known as phosphatidylinositol synthase (PIS), catalyzes the addition of a *myo*-inositol ring to a phospholipid backbone, cytidine diphosphate-diacylglycerol (CDP-DAG), to generate the base of all PIPs, phosphatidylinositol (PI) (Lykidis et al., 1997) (Fig 1A). This is the only protein currently known to perform this function in zebrafish (Thakur et al., 2011). CDIPT is a highly conserved integral membrane protein found on the cytoplasmic side of the endoplasmic reticulum (ER).

**Fig 1.**
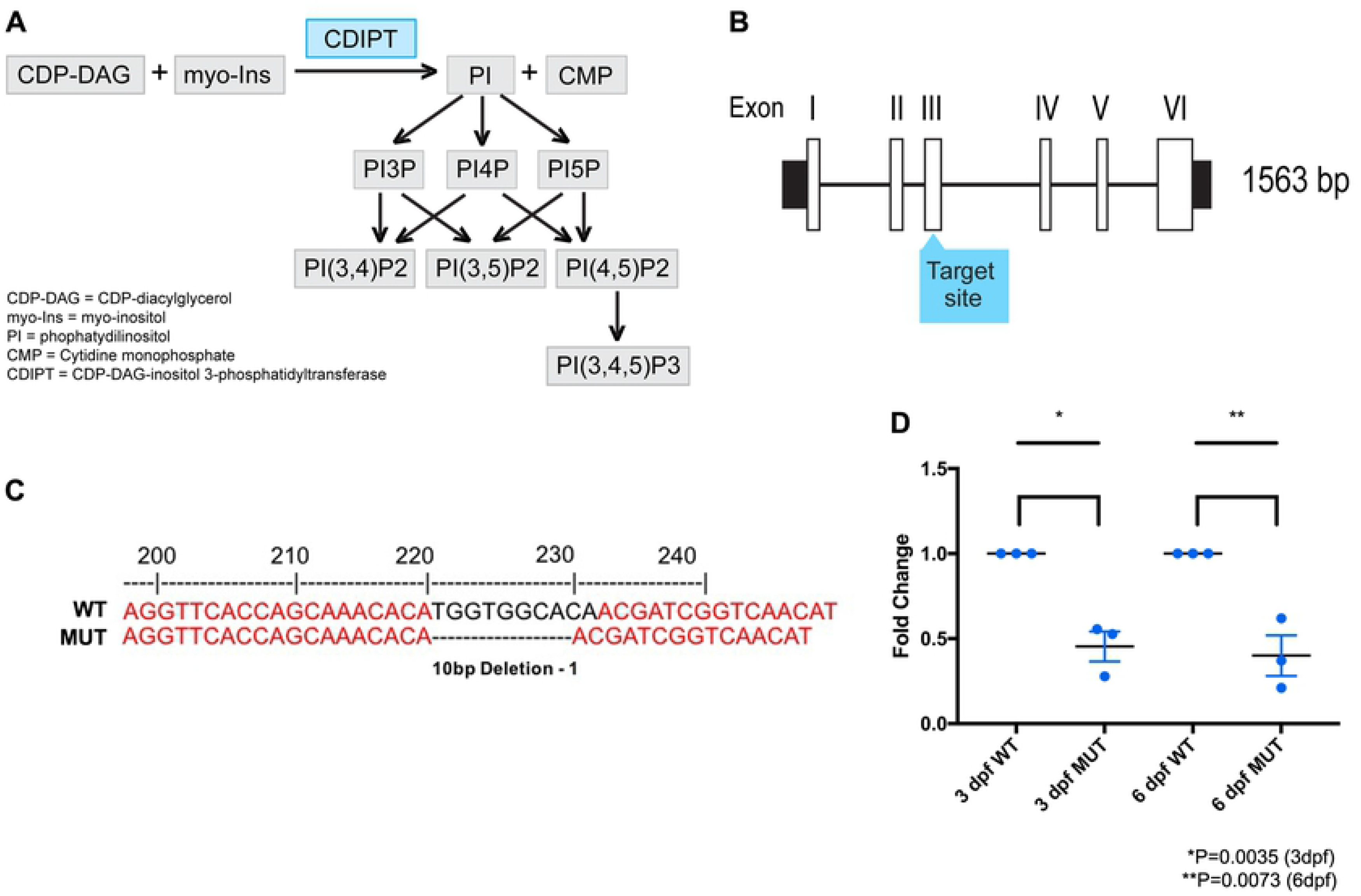
Development of a CRISPR/Cas9 *cdipt* mutant zebrafish. **A)** Schematic representation of phosphoinositide signaling pathway. CDIPT catalyzes the addition of the myo-inositol to the CDP-DAG to generate PI, which is the base precursor for all species of PIPs. **B)** Schematic representing exon organization of *cdipt.* Exon 3 was targeted by CRISPR/Cas9 gene editing. **C)** Sanger sequencing of wildtype (WT) and homozygous *cdipt* mutant (MUT) larvae showing a 10-bp deletion in exon 3 of *cdipt.* **D)** Fold change of mRNA levels between WT and MUT fish at both 3 dpf and 6 dpf. There is a significant change in *cdipt* mRNA levels between WT and MUT zebrafish at both 3 dpf (0.5-fold reduction; *p=0.0035) and 6 dpf (0.6-fold reduction; **p=0.0073). Each replicate is represented by a point, n = 30 per replicate; Student’s *t* test, 2-tailed. Error bars indicate SEM.

Previous study of a zebrafish *cdipt* mutant revealed a liver phenotype reminiscent of phenotypes seen in other models of PIP dysregulation (Thakur et al., 2011). This study, however, did not examine skeletal muscle. In the current study, we examine the skeletal muscle in a new *cdipt* mutant. We show that loss of CDIPT has no effect on early muscle development, suggesting that skeletal myogenesis does not require *de novo* PIP synthesis. Instead, CDIPT appears to be required for proper formation of the triad.

## Materials and methods

### Zebrafish maintenance

Zebrafish stocks were maintained at the Zebrafish Facility at the Hospital for Sick Children, Toronto, ON, Canada. All zebrafish procedures were performed in compliance with the Animals for Research Act of Ontario and the Guidelines of the Canadian Council on Animal Care.

### Generation of zebrafish *cdipt* mutants

A detailed procedure for CRISPR/Cas9 editing in zebrafish has been described previously (Ma et al., 2016). The *cdipt* target in this study was 5’-GGTTCACCAGCAAACACATGGTGG-3’ in exon 3. One-cell-stage AB WT embryos were injected with gRNA and Ca9 mRNA with a Picopump (World Precision Instruments). Potential founders (F_0_) were outcrossed to AB WT fish. Genomic DNA was isolated from single F_1_ embryos at 6 dpf and genotyped using high resolution melt (HRM) analysis. A *cdipt* sequence spanning the CRISPR/Cas9 target site was amplified with the following primers: F: 5’-AGCTGGAACAGAAAAGTGTAGGA-3’; and R: 5’-TAGGTACAAAATTTGGTGCAATG-3’. Carriers were identified and outcrossed ultimately to the F_3_ generation. In-cross progeny from the F_3_ and F4 generations were characterized in this study.

### Real-time PCR (qPCR)

RNA was extracted from 3 dpf and 6 dpf *cdipt* mutant zebrafish and their wildtype siblings using RNAeasy (Qiagen). RNA samples were reverse transcribed into cDNA using the iScript cDNA synthesis kit (BioRad). Primers were designed to result in a product spanning exons 4-6 of *cdipt:* F: 5’-ACCCCATTTTACGGCTGTACT-’3; and R: 5’-TACCTGGGGTTCTTCGATGT-’3. Products were amplified using Step-One-Plus Real-Time PCR System (Applied Biosystems). The zebrafish beta-actin gene, *actb1*, was used as an endogenous control.

### Birefringence

Tricaine-anaesthetised larvae were mounted in 3% methylcellulose on glass slides and imaged under polarized light on a dissecting microscope (Olympus SZX7).

### Skeletal myofiber preparations

Myofiber preparations of 6 dpf wildtype and *cdipt* mutant zebrafish were made following the protocol described previously (Horstick et al., 2013).

### Immunofluorescence staining

Immunostaining of myofiber preparations was performed as previously described (Horstick et al., 2013). Briefly, myofiber preparations were fixed with 4% PFA, for 20 min, permeabilized with PBST (0.3% TritonX in PBS), blocked for 1 hour with PBSTB (5% BSA in PBT) and incubated overnight at 4°C with primary antibodies. The following primary antibodies were used: mouse anti-DHPR (1:200; DHPRa1A; Abcam), mouse anti-α-Actinin (1:100; Sigma), mouse anti-RyR1 (1:100; 34C; DSHB), rabbit anti-Junctin (1:350; gift from Dulhunty lab), mouse anti-PI(3)P (1:100; Echelon Biosciences Inc.), mouse anti-PI(3,4)P2 (1:100; Echelon Biosciences Inc.), mouse anti-PI(4,5)P2 (1:100; Echelon Biosciences Inc.). Alexa Fluor-conjugated secondary antibodies were used at 1:1000 (Invitrogen). Rhodamine phalloidin (Phalloidin 555) was used to visualize filamentous actin (1:300, Molecular Probes). Preparations were mounted with ProLong Gold with DAPI (Invitrogen). Images were acquired with a Nikon Eclipse Ti laser scanning confocal using NIS Elements software (company, location) and only adjusted for brightness and contrast using Adobe Photoshop.

### Live confocal imaging

One-cell stage zebrafish embryos were injected with 10 pg of a cDNA construct containing a fluorescent protein attached to a PIP-binding protein domain [Bodipy-PI (Echelon Biosciences Inc)]; PLC-□-PH-GFP (Tobias Meyer Lab, Stanford University, CA) using a Picopump (World Precision Instruments). At 1 dpf, injected zebrafish were incubated in 0.2 mM phenylthiourea to prevent pigment formation. To image, zebrafish were screened for fluorescent myofibers on a macroscope (Zeiss Axio Zoom) and mounted in 1.5% low-melt agarose on a 3 cm glass-bottom petri dish. All confocal images were taken with a Nikon Eclipse Ti confocal microscope using a 40x oil-immersion lens.

### Transmission electron microscopy

Zebrafish clutches at 6 dpf were anaesthetised in tricaine and fixed in Karnovsky’s fixative overnight at 4°C. Samples were sent to the Advanced Bioimaging Center (Sickkids Peter Gilgan Centre for Research and Learning, Toronto) where larvae were processed. Briefly, larvae were rinsed in buffer, post-fixed in 1% osmium tetroxide in buffer, dehydrated, and embedded in Quetol-Spurr resin. Following this, 70 nm sections thick were cut with a Leica UC7 ultramicrotome, stained with uranyl acetate and lead citrate, and viewed either with an FEI Tecnai 20 transmission electron microscope (Technai, Oregon, USA) (Bioimaging Facility at The Hospital for Sick Children, Toronto) or with a JEOL JEM 1200EX TEM (JEOL, Massachusetts, USA) (Electron Microscopy Facility at the Laboratory of Pathology, The Hospital for Sick Children, Toronto). Images were obtained using Gatan Digital Micrograph acquisition software or AmtV542, and were manipulated only for brightness and contrast using Adobe Photoshop.

### Immuno-electron microscopy

6 dpf zebrafish embryos were anaesthetised in tricaine and fixed for 2h at room temperature followed by overnight fixation at 4°C in 4% PFA, 0.1% glutaraldehyde in 0.1M sodium cacodylate buffer with 0.2M sucrose. Samples were rinsed in 0.1M sodium cacodylate buffer and dehydrated in ethanol series (70% ethanol for 1h at 4°C; 90% ethanol for 1h at 20°C; 100% ethanol for 1h at −20°C, twice). Samples were then embedded in 50/50 LR White resin/ethanol for 1h at −20°C, followed by 70/30 LR White resin/ethanol for 1h at −20°C and 100% LR White resin for 1h at −20°C. Samples were then left overnight at −20°C in 100% LR White resin. Embryos were then placed in capsules filled with LR White resin mixed with benzoin methyl ether (0.1 g in 100 ml LR White), sealed, and placed in the oven for polymerization at 65°C for at least 72h.

70 nm ultrathin sections were cut with a Leica UC7 ultramicrotome and placed on formvar-coated grids, which were then processed for gold labelled immunostaining. Grids were treated with 0.15M glycine in PBS for 15 minutes, rinsed with PBS, followed by Aurion blocking solution (Aurion, The Netherlands) for 15 minutes. Primary antibodies were diluted in 0.1% BSA-c (Aurion, The Netherlands) at the following concentrations: 1:25 mouse anti-PI(4,5)P_2_; 1:10 mouse anti-PI(3,4)P_2_; 1:10 mouse anti-PI(3)P (Echelon Biosciences, Inc.). Samples were incubated with primary antibodies for 1h at room temperature. After rinsing the samples with PBS 5 x 5 minutes, these were incubated with 10nm gold-conjugated goat anti-mouse secondary antibody (1:10; Electron Microscope Sciences, Hatfield, PA) for 1h at room temperature. Samples were then rinsed 5 x 5 minutes with PBS, treated with 2% glutaraldehyde (in PBS) for 5 minutes, rinsed 5 x 5 minutes with distilled water and air dried. Gold immunolabelled samples were counter-stained with uranyl acetate and lead citrate and viewed with a JEOL JEM 1200EX TEM (JEOL, Massachusetts, USA) (Electron Microscopy Facility at the Laboratory of Pathology, The Hospital for Sick Children, Toronto). Images were obtained using AmtV542 software, and were manipulated only for brightness and contrast using Adobe Photoshop.

### Triad size measurement

To determine total triad area (A1+A2+A3), the following features were measured: area of T-tubule (A1), areas of the two terminal cisternae (A2, A3), the distance between the membranes of the two terminal cisternae (D1) and the width of the gap between the membrane of the terminal cisternae and the T-tubule membrane (*). Measurements were done using the open source software Image J. Data and statistical analyses were performed using GraphPad Prism 8 (GraphPad Software Inc., San Diego, CA).

### Swim test and photoactivation assay

All motor behaviour analysis was performed using Zebrabox software (Viewpoint, France) as previously described (Sabha et al., 2016). To perform the photoactivation assay, zebrafish were incubated for 5 min at 28.5°C with optovin 6b8 (ID 5705191l; ChemBridge), an optovin analog (Kokel et al., 2013). Optovin is a reversible TRPA1 ligand that elicits motor excitation following exposure to light. After incubation, the Zebrabox platform monitored larvae for 20 second cycles over 10 minutes. Parameters were set to capture 5 seconds of exposure to white light to elicit ambulatory movement, followed by 15 seconds of recovery behaviour in the dark. This was repeated 30 times, to get a total experiment time of 10 minutes. The average speed traveled during the 20 second cycle was used to compare groups (i.e., *cdipt* mutants vs. WT siblings). To perform the spontaneous swim assay, zebrafish in system water were followed for 1 hour. Data were analyzed using statistics software (GraphPad Prism).

### Morpholino studies

For knockdown of maternal *cdipt,* the following ATG-targeting MO was designed: 5’-CCGAGAGTTTCTTTCTTTGGACGGA-’3 (GeneTools LLC). An MO designed to a random sequence (5’-CCTCTTACCTCAGTTACAATTTATA-3’) with no homology by Basic Local Alignment Search Tool (BLAST) analysis in the zebrafish genome was used as a control (GeneTools LLC). Fertilized eggs were collected after timed matings of adult zebrafish and injected at the 1-cell stage using a Picopump (World Precision Instruments). Embryos were injected with concentrations ranging from 0.15 – 0.5 mM in a volume of 1 nl.

## Results

### Developing a new *cdipt* mutant zebrafish line

Exon 3 of the *cdipt* gene was targeted using the CRISPR/Cas9 system (Fig 1B). A 10 bp deletion allele, hereafter referred to as *cdipt* mutant when present in homozygosity, was identified after Sanger sequencing (Fig 1C). Due to lack of commercially available antibodies against zebrafish CDIPT, we performed real-time PCR (qPCR) on total RNA from whole embryos to confirm that *cdipt* transcript is reduced by this mutation. There was a significant difference in *cdipt* mRNA levels between wildtype (WT) and mutants at both 3 dpf and 6 dpf (Fig 1D) (P < 0.05, n = 30). This suggests that mutant *cdipt* mRNA transcripts are being directed to the nonsense-mediated decay pathway, and is consistent with this mutation being a loss of expression and function allele.

### *cdipt* zebrafish exhibit morphological and gastrointestinal system abnormalities

Homozygous *cdipt* mutant fish appeared phenotypically normal until 5 dpf, when gastrointestinal system abnormalities are visible with bright-field microscopy. The mutant phenotype is fully penetrant at 6 dpf and includes a dark, globular liver and small intestine, partial deterioration of the ventral fin (folds, incisions, missing areas), tissue degradation around the cloaca, and abnormal jaw structure (Fig 2A and S1 Fig). The gastrointestinal features are reminiscent of the mutant *cdipt^hi559/hi559^* phenotype, and have already been well characterized (Thakur et al., 2011).

**Fig 2.**
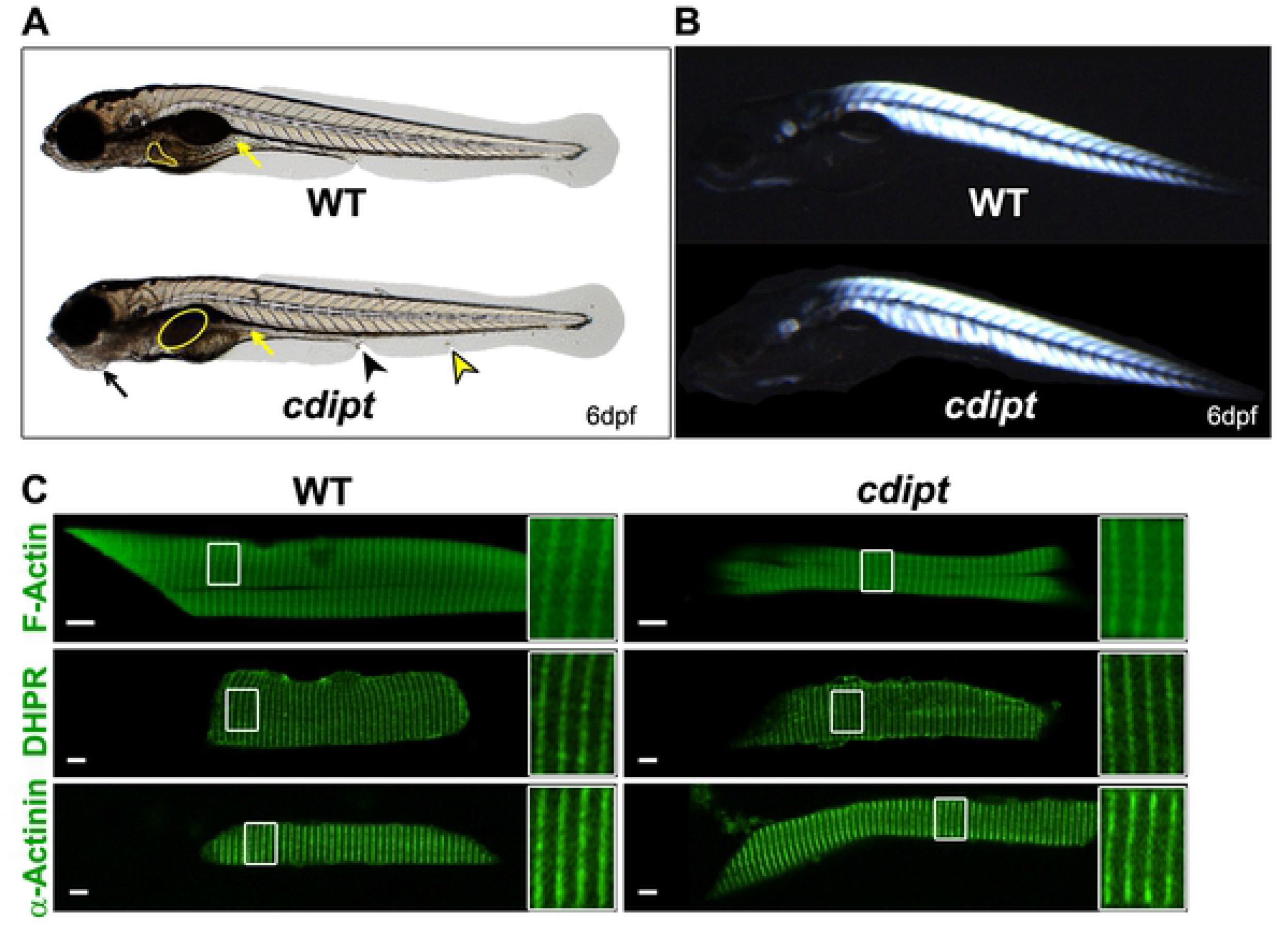
Characterization of the *cdipt* mutant phenotype at 6dpf. **A)** *cdipt* mutant zebrafish exhibit a gastrointestinal phenotype with a dark, globular and oversized liver (yellow outline) and a small intestine (yellow arrows), abnormal jaw structure (black arrows), tissue degradation around the cloaca (black arrowhead), and defective ventral fin (yellow arrowhead). **B)** Representative image of *cdipt* mutants at 6dpf, showing normal birefringence pattern indistinguishable from WT siblings, indicative of normal sarcomere organization. **C)** Confocal micrographs showing localization by indirect immunoflourescence of actin (upper panels), DHPR (middle panels) and α-actinin (bottom panels) in the skeletal myofibers. There is no noticeable difference in the localization of these proteins between WT (left column) and *cdipt* mutant (right column). Insets represent high magnification of areas surrounded by white rectangles. Scale bars = 5 μm.

### *cdipt* zebrafish have generally normal muscle structure

Gross morphology of *cdipt* zebrafish muscle was investigated using birefringence. Birefringence uses polarized light to assess muscle integrity. Organized skeletal muscle will appear bright amidst a dark background when visualized between two polarized light filters, whereas disorganized muscle exhibits degenerative dark patches and an overall decrease in brightness in some myotomes. Based on birefringence analysis, *cdipt* zebrafish have normal muscle integrity and sarcomere organization at all ages examined (Fig 2B).

We next studied the localization of several sarcomeric proteins, such as actin, myosin (contractile proteins), dihydropyridine receptor (DHPR), ryanodine receptor type 1 (RyR1) and junctin (markers for triads), laminin and dystrophin (markers for myotendinous junctions), and α-actinin (a Z-line marker). Immunostaining with antibodies against these proteins on myofibers isolated from 6dpf zebrafish showed no differences in localization between WT and *cdipt* mutants (Fig 2C and S2 Fig), indicating no qualitative defects in the formation and organization of key muscle structures.

We next studied the ultrastructure of muscle, given that abnormal triad formation is a hallmark of many PIP-related myopathies and may not be appreciated by light microscopy. *cdipt* larvae and their WT siblings were thus processed at 6 dpf for transmission electron microscopy. Electron micrographs revealed no major abnormalities in triad structure in *cdipt* mutants (Fig 3A,B). To better characterize the triads we measured the area of T-tubules (A1), the areas of the two terminal cisternae (A2, A3), total triad area (A1+A2+A3), the distance between the membranes of the two terminal cisternae of the sarcoplasmic reticulum (D1), and the width of the gap between the T-tubule and each of the two terminal cisternae (*), where the junctional feet corresponding to the ryanodine receptor-dihydhropyridine receptor complex are found (Al-Qusairi and Laporte, 2011) (Fig 3C). There was no significant difference in the total area of the triad (A1+A2+A3) (Fig 3D). However, the T-tubule area (A1) was qualitatively slightly smaller in the *cdipt* mutant as compared to WT, and the distance between terminal cisternae at maximum distance (D1) and the gap width (*) were quantitatively and significantly smaller in *cdipt* mutants (Fig 3D).

**Fig 3.**
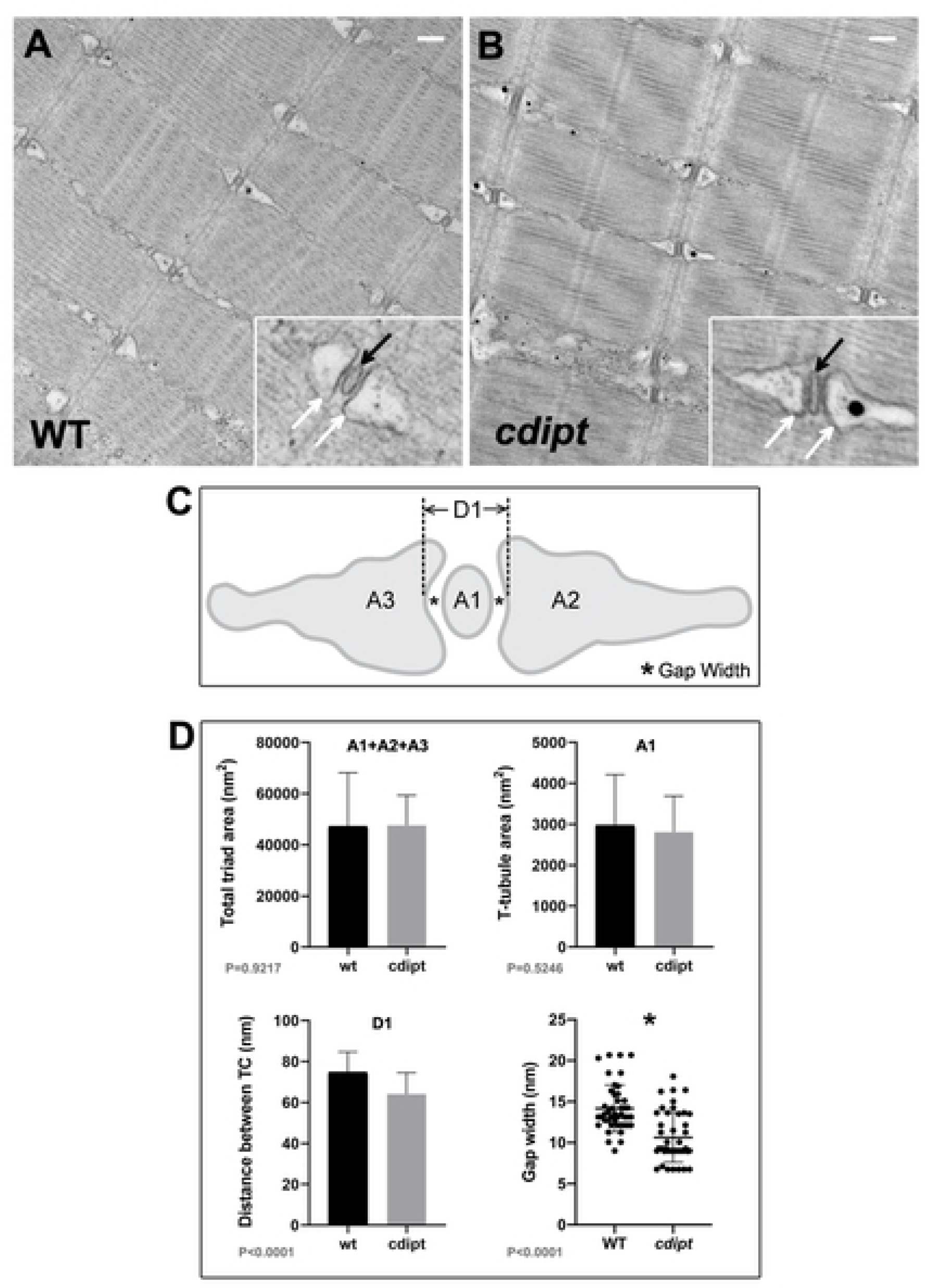
Skeletal muscle ultrastructure. **A-B)** Transmission electron micrographs show normal skeletal muscle ultrastructure in *cdipt* larvae. T-tubules (insets; black arrow) are apposed by terminal cisternae of sarcoplasmic reticulum (insets; white arrow). **C)** Diagram illustrating triad structure and features used for measurements: A1 = T-tubule area; A2 and A3 = terminal cisternae (TC) areas; D1 = maximum distance between TCs; * = gap width (distance between TC membrane and T-tubule membrane). **D)** There is no significant difference in the triad area between WT and *cdipt* mutant larvae (A1+A2+A3 graph) (n = 36, p = 0.9217). The T-tubule area (A1 graph) is qualitatively slightly smaller in the *cdipt* mutant than in WT (n=36; P=0.5246), whereas the distance between cisternae at maximum distance (D1 graph) (n = 36, p < 0.0001) and the gap width (* graph) (n = 44, p < 0.0001) are significantly smaller in *cdipt* mutants than in WT. Scale bars = 200 nm.

### *cdipt* zebrafish have abnormal motor behaviour as compared to wildtype siblings

Given the subtle but significant change observed in the appearance of the triad, we wanted to determine if there was any alteration in muscle function. To assess muscle function, we performed a spontaneous swim test assay and a routine photoactivation movement assay previously utilized by our lab (Sabha et al., 2016). The latter involved incubating zebrafish larvae with a molecule called optovin 6b8, which when exposed to white light, will activate zebrafish muscle through a reflex arc. If muscle function is impaired, mutants will have reduced movement when compared to WT. We did thirty rounds of 20 second-activation periods to assess both the speed of movement and muscle fatigue. The average speed of movement was significantly lower in *cdipt* mutant zebrafish both in the spontaneous swim test (Fig 4A,B) and in their response to optovin (Fig 4C,D). In addition, *cdipt* mutants spent less time moving (S3A Fig) and covered shorter distances than their WT siblings (S3B Fig). The rate of fatigue, however, was similar in the *cdipt* mutant and WT zebrafish (Fig 4D).

**Fig 4.**
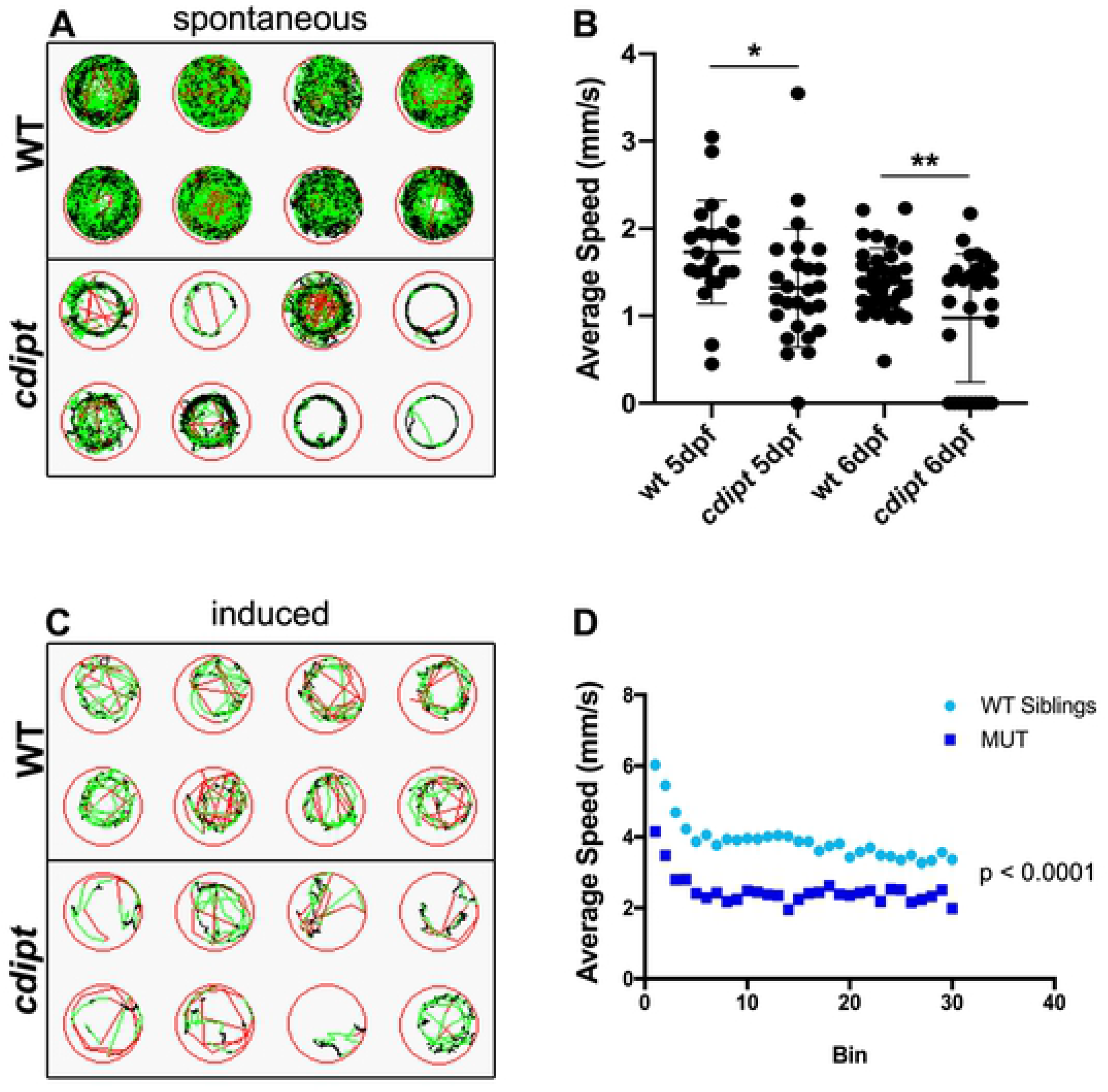
*cdipt* mutants have significantly impaired motor function compared to their wildtype siblings. **A)** Spontaneous swim movement was assessed by tracking 5-days or 6-days old zebrafish larvae over 1 hour. Representative examples of tracking plots of individual larvae movement. Black represents slow movement (<5 mm/s), green represents average speed (5-20 mm/s), and red represents fast movement (>20 mm/s). **B)** The *cdipt* mutant larvae are significantly slower than their WT siblings, both at 5dpf (WT n = 22, *cdipt* n = 26, p = 0.0318) and 6dpf (WT n = 36, *cdipt* n = 28, p = 0.0036). **C)** Involuntary motor function was assessed using an optovin-stimulated movement assay in response to pulses of light. Representative examples of tracking plots of individual larvae showing movement over 20 seconds, involving 5 seconds of white light exposure followed by 15 seconds of darkness. **D)** There is a significant difference between the average speed travelled by WT zebrafish compared to *cdipt* mutant zebrafish (n = 18 and 14, respectively; p < 0.0001, Student’s *t* test, 2-tailed). WT and *cdipt* mutant zebrafish plateau at the same rate (n = 7, p=0.3487).

### *cdipt* zebrafish do not show changes in the localization of PIPs

Given that CDIPT is the rate-limiting enzyme for PI synthesis (the precursor for all PIPs), we looked at the expression and localization of several species of PIPs in myofibers. We specifically investigated PI3P, PI(3,4)P2 and PI(4,5)P2 localization. PI(4,5)P2 and PI(3,4)P2 are found mostly at the plasma membrane, whereas PI3P is mostly found on endosomes (Viaud et al., 2016). Immunofluorescence staining with anti-PIP antibodies revealed no significant differences in the localization of PI(4,5)P2, PI3P, PI(3,4)P2 (Fig 5). To further investigate PI(4,5)P2 localization in *cdipt* larvae *in vivo,* a fluorescent marker for PI(4,5)P2 (PLCδPH-GFP) was injected into 1-cell stage *cdipt* embryos. At 6 dpf, PI(4,5)P2 appeared to properly localize to the plasma membrane (Fig 5A,A’). These results were further supported by immunoelectron microscopy studies. Nanogold-labelled antibodies against PI3P, PI(3,4)P2 and PI(4,5)P2 localized at the triad and its vicinity and showed similar localization pattern in WT and *cdipt* mutant embryos (Fig 6).

**Fig 5.**
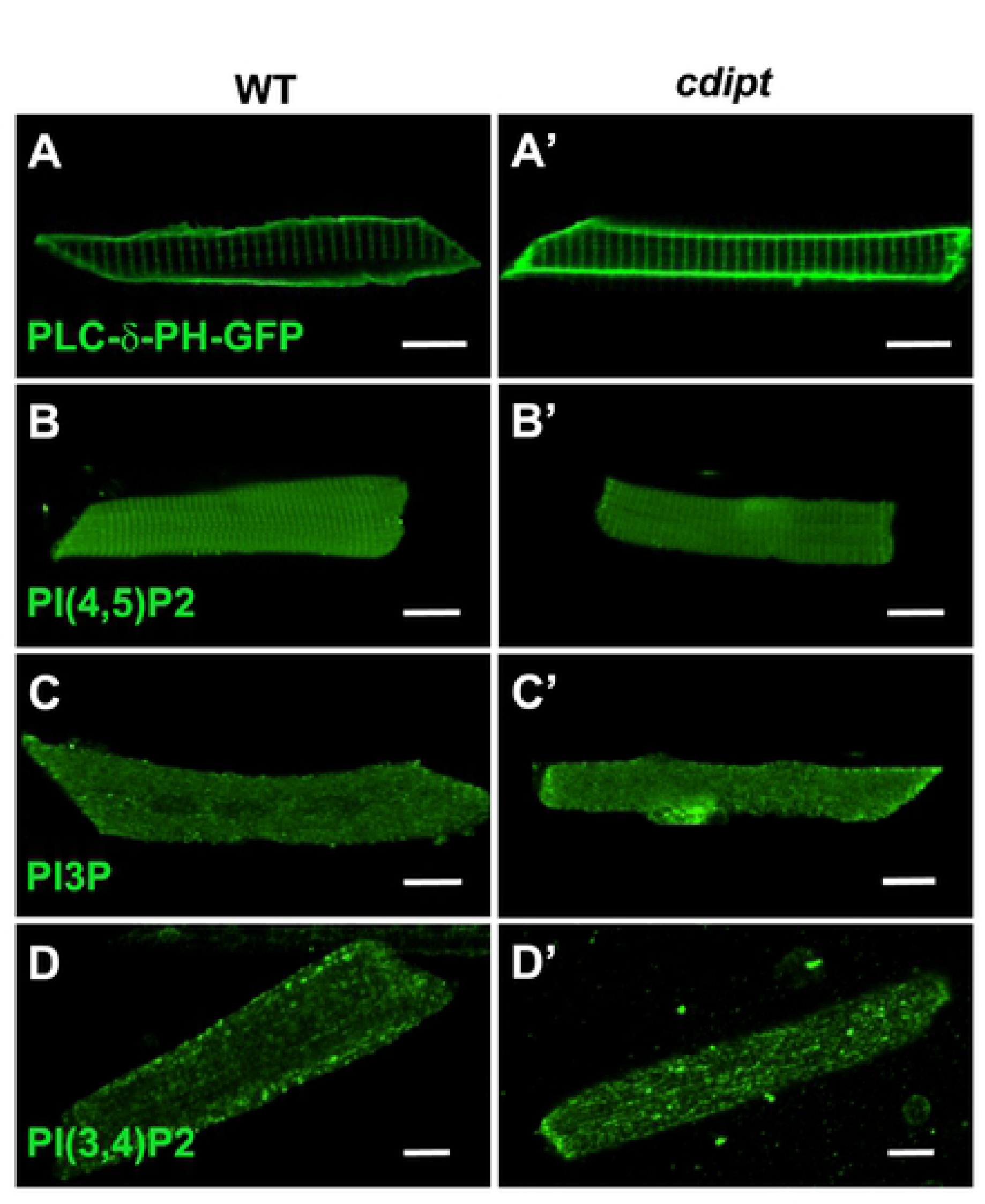
Localization of PIPs by immunofluorescence in wildtype and *cdipt* mutant zebrafish. Confocal micrographs showing localization of PIPs is not affected in early larval development of *cdipt* mutants. **(A, A’)** visualization of skeletal muscle from live embryos injected with PLCδPH-GFP, a marker for PI(4,5)P_2_. There was no obvious difference in expression between wild type (WT) and *cdipt* mutant embryos. **(B-D, B’-D’)** Immunostaining with PIP antibodies of myofibers isolated from WT and cdipt mutants. Localization of PI(4,5)P_2_ **(B, B’)**, PI3P **(C, C’)** and PI(3,4)P_2_ **(D, D’)** is similar in wildtype and *cdipt* zebrafish. Scale bars = 10μm.

**Fig 6.**
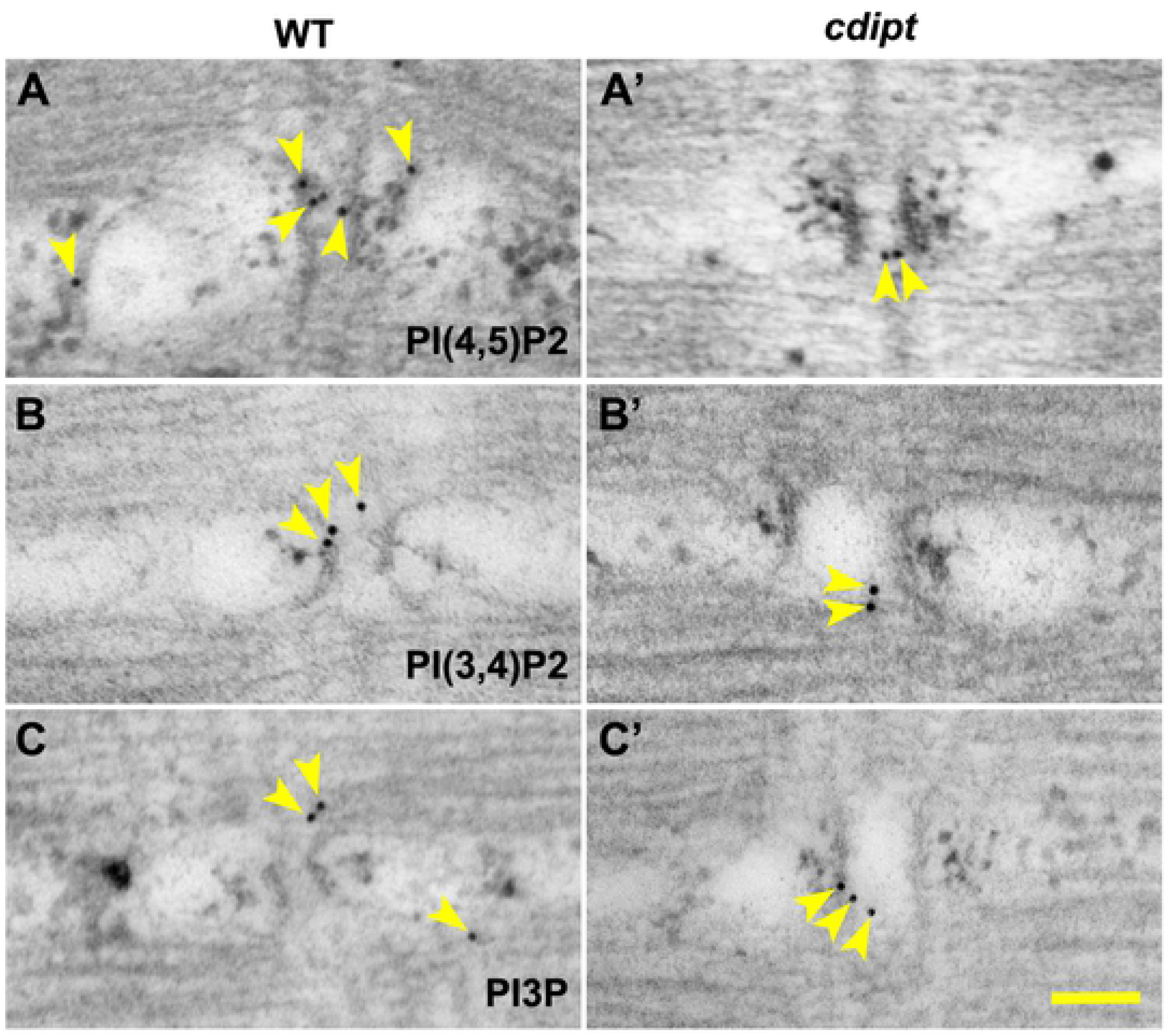
Localization of PIPs by immunoelectron microscopy in wildtype and *cdipt* mutant zebrafish. Transmission immunoelectron micrographs showing localization of nanogold-labelled antibodies against **(A, A’)** PI(4,5)P_2_, **(B, B’)** PI(3,4)P_2_ and PI3P **(C, C’)** (yellow arrowheads) at the skeletal muscle triad. There is no difference in localization of these antibodies between WT and *cdipt* mutant embryos. Scale bar = 100nm.

### Maternal *cdipt* mRNA and/or PI are sufficient for normal muscle development

Given the importance of CDIPT in generating a precursor to all PIP species, it is surprising that there is no developmental phenotype in skeletal muscle. However, a previously published lipidomic analysis of the early zebrafish yolk found that PI is already present at 0 hpf (Fraher et al., 2016) and previous results on *cdipt^hi559/hi559^* zebrafish (Thakur et al., 2011) and our qPCR results (see Fig 1D) show there is maternal *cdipt* mRNA expression in early stages of the zebrafish embryo before zygotic gene expression is turned on.

To prevent production of CDIPT protein from maternal mRNA, we injected zebrafish embryos at one-cell stage with a translation blocking morpholino (ATG-MO). Injection of *cdipt* ATG-MO at both 0.3 mM and 0.5 mM caused increased levels of embryo death (S4A Fig), suggesting that blocking of maternal *cdipt* mRNA translation is broadly detrimental for embryogenesis. While we were not able to study muscle in the majority of *cdipt* ATG morphants due to early lethality, some morphants did survive beyond the first day post fertilization (S4B Fig). In those morphants, the skeletal muscle development was not obviously affected.

To investigate whether maternally deposited PI in the yolk can be delivered to developing skeletal muscle, we injected BODIPY-labelled PI into yolk at the one-cell stage. Fluorescence was tracked with a confocal microscope over several days. By 1 dpf, the fluorescent probes appeared in the skeletal muscle compartment (S5 Fig). Fluorescence was not detectable at 2 days post-injection and later. These results suggest that PI present in the yolk at early stages can be delivered to skeletal muscle. Taken together, our data suggest that the presence of maternally deposited mRNA in the cell and/or PI in the yolk fulfill the early developmental requirements for CDIPT and for PI, which is consistent with the lack of a phenotype until the yolk is depleted at 5 dpf. This also suggests that once PI and its PIP derivatives are generated, they likely persist as a stable pool in skeletal muscle.

## Discussion

To investigate the role of PIPs in muscle development, we developed and characterized a new *cdipt* mutant zebrafish. This mutant showed defects in fin morphology and aberrant swimming behaviour, in addition to the gastrointestinal defects previously reported in another *cdipt* mutant *[cdipt^hi559/hi559^*, (Thakur et al., 2011)].

The purpose of generating this model was to determine how the potential loss of all seven PIP species would affect muscle development. We expected that loss of CDIPT and the subsequent depletion of PI would have severe effects on skeletal muscle; however, mutant *cdipt* larvae showed only minimal abnormalities in muscle structure and overall muscle function. We hypothesize that this modest phenotype is the result of PI deposited in the muscle during embryogenesis that then provides sufficient substrate for generation and maintenance of PIP species at subsequent developmental stages.

Given the importance of CDIPT in generating a precursor to all PIP species, it is surprising that there is no significant adverse phenotype in muscle. The lack of widespread abnormalities may be because *cdipt* mRNA and PIs are maternally deposited into zebrafish yolk. Depletion of maternal mRNA using a translation-blocking morpholino resulted in increased mortality in the zebrafish, consistent with a role for CDIPT and de novo PI synthesis in embryogenesis. However, the impact of the translation-blocking morpholino did not seem to affect skeletal muscle development in the surviving injected embryos, suggesting that maternally deposited CDIPT does not play a role in myogenesis. There is likely also an important contribution to total embryo PI from maternal deposition of this precursor lipid in the yolk. In order to remove this potential confounder to the assessment of CDIPT function in skeletal muscle, we would need to prevent the deposition of PI into the yolk, or develop a method of depleting yolk PI without disrupting other essential nutrients contained within the yolk. Currently, there are no technologies that would allow us to complete either of those experiments. We attempted to use direct lipase injection into the yolk in order to deplete it, but this resulted in embryonic lethality prior to myogenesis.

The one part of the muscle where we did observe abnormalities was the triad. We showed by immunofluorescence that several species of PIPs localize to the sarcomere. Moreover, our novel immunoelectron microscopy studies showed these molecules localize to the triad. To our knowledge, this is the first report on ultrastructural localization of PI3P, PI(3,4)P2 and PI(4,5)P2 at the triad in a vertebrate model, as previous studies focused on culture cells (Watt et al., 2002) (Tabellini et al., 2003) (Mayhew et al., 2004) (Wegner et al., 2014) (Pastorek et al., 2016). Our data add to the growing evidence showing the importance of PIP metabolism in the development and maintenance of this key muscle substructure. Work with mammalian myocytes in culture has implicated PIP2 (via its binding to BIN1) in the formation of the T-tubule. While we did not see an overall decrease in PIP2 levels, nor in its localization, it is tempting to speculate that the loss of CDIPT sufficiently impacted PIP synthesis enough to result in mild by critical reductions in PIP2 that were enough to alter triad formation. Future studies will be required to fully explore this relationship.

Of note, phenotypic abnormalities in *cdipt* mutants do not appear until 5 dpf, shortly after the yolk has been depleted. The most prominent of these is the digestive system phenotype, likely reflecting a requirement for *de novo* PIP synthesis in this organ system, since pools of PI are locally made and used almost immediately after synthesis (Varnai and Balla, 2006). However, because skeletal muscle has no phenotype at 5 dpf and previous data has shown that *cdipt* mRNA is not present in skeletal muscle after 5 dpf (Varnai and Balla, 2006), it is possible that maturing skeletal muscle does not require *de novo* PIP synthesis. Instead, perhaps PIPs are maintained in pools that can fluctuate between the different species when needed. Alternatively, a requirement for PIP synthesis in muscle may not manifest in the window of time after yolk depletion and before mutant death, but may develop as the muscle continues to grow and mature. Muscle specific targeting of *cdipt* would be helpful in the future to distinguish between these possibilities.

Of note, the most obvious phenotypes in the *cdipt* mutants are in the liver, gastrointestinal system, and the fin. Interestingly, these phenotypes of the *cdipt* mutants are also visible in *mtm1* mutant zebrafish (Sabha et al., 2016). The fact that two mutated PIP-related genes cause similar defects suggest that PIP metabolism must be tightly regulated in these tissues in zebrafish development, and that there is an increased requirement for de novo synthesis and homeostatic balance.

## Acknowledgements

The authors gratefully thank Jonathan Volpatti, Yukari Endo and Mo Zhao (Dowling Lab) for useful discussions and insightful suggestions; Evangelina Aristegui for technical support; Scott Knox, Alejandro Salazar, and Elyjah Schimmens for zebrafish care (SickKids Zebrafish Facility); Paul Paroutis and Kimberley Lau for technical support (SickKids Imaging Facility); and Doug Holmyard, Ali Darbandi and William Martin (SickKids Nanoscale Biomedical Imaging Facility) for help with TEM and immunoEM sample preparation. The authors also thank the following people for generously providing reagents: Angela Dulhunty (John Curtin School of Medical Research, Australian National University, Canberra, Australia) for anti-Junctin antibody; Tobias Meyer Lab (Stanford University, CA) for the PLC-δPH-GFP construct; Sergio Grinstein and Julie Brill (The Hospital for Sick Children, Toronto, Canada) for PIP constructs and for helpful discussion.

## Supplemental material

**Fig. S1. Phenotypic variations in cdipt mutants. A-B)** Examples of fin degeneration (yellow arrowheads), oversized liver (yellow outline), and abnormal jaw structure (black arrows) in *cdipt* mutant zebrafish. **C)** Many *cdipt* mutants have partially folded ventral fin.

**Fig. S2. Localization of triad-associated proteins in the muscle**. Confocal micrographs showing localization by indirect immunofluorescence of RyR1 (top panels) and Junctin (bottom panels) in skeletal myofibers. There is no noticeable difference in localization of these proteins in WT (left panels) and *cdipt* mutant (right panels). Scale bars = 10 μm.

**Fig. S3. cdipt mutants have impaired motor function**. **A)** *Cdipt* mutant zebrafish spend significantly less time swimming compared to their WT siblings, both at 5dpf (WT n = 22, *cdipt* n = 26, p = 0.0359) and 6dpf (WT n = 36, *cdipt* n = 28, p = 0.0209). **B)** *Cdipt* mutant zebrafish travel significantly shorter distances compared to their WT siblings at 5dpf (WT n = 22, *cdipt* n = 26, p = 0.0121) whereas at 6dpf the travelled distances are not significantly different (WT n = 36, *cdipt* n = 28, p = 0.1015).

**Fig. S4. Blocking maternal cdipt mRNA translation is detrimental for embryogenesis**. **A)** Embryos injected with ATG-MO (n=115) show significantly higher mortality rates than those injected with Ctrl-MO (n=117). **B)** Surviving ATG-MO-injected *cdipt* embryos have a normal birefringence pattern indistinguishable from their WT siblings.

**Fig. S5. Maternally deposited PI in the yolk is transported to the muscle.** Zebrafish larvae at 1 dpf after injection of BODIPY-PI into yolk at the 1-cell stage (arrows indicate accumulation of fluorescently-labeled PI in the muscle).

